# Humans Exploit the Trade-Off Between Lateral Stability and Manoeuvrability During Walking

**DOI:** 10.1101/2023.06.06.543488

**Authors:** Rucha Kulkarni, Francis M. Grover, Anna Shafer, Xenia Schmitz, Keith E. Gordon

**Affiliations:** Department of Biomedical Engineering, Northwestern University, McCormick School of Engineering, Evanston IL 60208; Physical Therapy and Human Movement Sciences, Northwestern University, Feinberg School of Medicine, Chicago, IL 60611; Research Service, Edward Hines Jr. VA Hospital, Hines, IL 60141

**Keywords:** Balance, gait, locomotion, reaction time, margin of stability

## Abstract

People use the mechanical interplay between stability and manoeuvrability to successfully walk. During single limb support, body states (position and velocity) that increase lateral stability will inherently resist lateral manoeuvres, decrease medial stability, and facilitate medial manoeuvres. Although not well understood, people can make behavioural decisions exploiting this relationship in anticipation of perturbations or direction changes. To characterize the behavioural component of the stability-manoeuvrability relationship, twenty-four participants performed many repetitions of a discrete stepping task involving mid-trial reactive manoeuvres (medial or lateral direction) in a Baseline (no external perturbations) and Perturbed (random mediolateral perturbations applied to their pelvis) environment. We hypothesized people would make systematic changes in lateral stability dependent on both environment (increasing lateral stability in the Perturbed environment) and anticipated manoeuvre direction (reducing lateral stability to facilitate lateral manoeuvres). Participants increased lateral stability in the Perturbed environment, coinciding with an increase in manoeuvre reaction time for laterally but not medially directed manoeuvres. Moreover, we observed lower lateral stability in both environments when people anticipated making a lateral manoeuvre when compared to medial manoeuvres. These results support the hypothesis that people behaviourally exploit the mechanical relationship between lateral stability and manoeuvrability depending on walk task goals and external environment.

## 1.0 Introduction

From aircrafts (1,2) to robotics (1,3) to animals (4–7), substantial research has studied how body design influences movement stability and manoeuvrability. A major finding is that characteristics that improve stability limit manoeuvrability and vice versa. Broadly speaking, in animal locomotion, stability is defined as a resistance to, and recovery from disturbances to an intended movement trajectory, and manoeuvrability, as the magnitude of the control signal required to change direction (6). This inverse relationship implies that stable body designs that resist external perturbations will also inherently resist volitional efforts to change directions (8). To understand how this stability-manoeuvrability trade-off impacts human walking, it is important to recognize that some gait characteristics are flexible. For example, people have the capacity to change their body dynamics and generate external forces that can change their state along the stability-manoeuvrability relationship on an ongoing basis during movement. Thus, behavioural decisions can impact the mechanical interplay between stability and manoeuvrability. How people choose walking behaviours that change their state along the stability-manoeuvrability relationship in situations that necessitate gait that is both stable and manoeuvrable is not well understood.

Real-world situations, such as walking quickly through a moving crowd of people, are challenging because they require gait patterns that are at times both stable (resistant to perturbations) and manoeuvrable (responsive to volitional efforts to quickly and accurately change direction). To address these needs, people can make behavioural changes to their gait patterns to bias the stability-manoeuvrability relationship in a manner consistent with their anticipated external threat to stability and anticipated need to manoeuvre. For example, many studies have identified that people will adapt gait patterns that increase their lateral stability in response to ongoing or potential threats of lateral perturbations (9–11). Other studies have observed that people will adapt gait patterns that reduce their resistance to lateral perturbations (less stable) in anticipation of an upcoming lateral manoeuvre (12,13).

Collectively this prior research indicates that behavioural leveraging of gait patterns is a common strategy people use to manipulate their state along the stability-manoeuvrability relationship in anticipation of their external environment and task goals. While these behavioural decisions have been observed in isolated situations that consider either stability or manoeuvrability, we do not know how people select gait patterns in complex walking environments that require both stability and manoeuvrability.

Thus, the objective of this study was to quantify the underlying behavioural modifications people make to their gait patterns when walking in complex situations that require both stability and manoeuvrability. We had people perform many trials of a discrete walking task—a rapid series of three steps that moved a person from a static standing position to a specified end target location. The discrete trial design was used to control initial conditions (body position and velocity) known to affect resistance to external perturbations (14). During the series of discrete walking trials, we systematically introduced unpredictable external perturbations that could challenge stability. The goal of these external perturbations was to create an environment where people would anticipate a threat to their lateral stability. In isolation, the response to such an environment is to adapt gait patterns that increase lateral stability (11). To study the interactions between lateral stability and manoeuvrability, we also required participants, at unexpected moments, to reactively produce either laterally or medially directed manoeuvres. The goal of the irregular but reoccurring manoeuvres was to create a task where people would anticipate having to change their walking trajectory in a known direction. In isolation, the response to an anticipated manoeuvre task is adapt gait patterns that decrease stability in the anticipated direction of movement (12,13). We then evaluated changes in gait stability and manoeuvrability. Specifically, we compared how people modulated an instantaneous measure of lateral stability and how well they performed manoeuvres during each trial in response to environmental threat (Baseline environment: no external perturbations vs. Perturbation environment: unpredictable external perturbations) and anticipated manoeuvre direction (lateral vs. medial manoeuvres).

The quantification of gait stability is a critical component of this study. Various methods have been used to assess gait stability - depending on the biomechanical, dynamical, and task-relevant contexts. Dynamic measures of stability (e.g., Lyapunov exponents, Floquet multipliers, detrended fluctuation analysis) have been popular tools to estimate variability (or lack thereof) in steady-state gait over time (15–17). Such measures assess the divergence (Lyapunov exponent), or the rate of divergence (maximum Floquet multiplier), or the persistence of fluctuations (DFA) of stability in the human system from one state to the next and can test whether a period of steady-state gait is stable. However, due to the assumption of steady-state continuous walking, these measures require a high number of strides (∼ 200) to provide a valuable estimate of stability, and therefore are not suited to assess stability of discrete steps (15,18). In contrast, some measures, derived from biomechanics, have assessed the momentary lateral stability of the whole body centre of mass (COM) within the base of support (15,19,20). Moreover, measures have combined dynamic and biomechanical aspects to estimate the future state (21) or the probability (22) of gait instability based on a prior series of gait cycles.

Our study was focused on single limb support of an isolated step during discrete walking manoeuvres. We therefore selected Hof’s biomechanical approach (19,20), that uses a simple inverted pendulum model of walking, to gain insight into the momentary mechanical relationship between frontal-plane stability and manoeuvrability. In this model, a margin of stability (MOS) is calculated as the distance between the extrapolated centre of mass (XCOM)—a metric accounting for both COM position and velocity—and the base of support (BOS); the border of the stance foot aligning with the direction of COM motion (i.e., lateral border during lateral motion) (19). During walking, the impulse needed to move the XCOM beyond the BOS is proportional to the MOS (19). When the XCOM position exceeds the BOS, the system is no longer passively stable (as gravity will accelerate the COM further beyond the BOS) and will require a corrective action (such as a cross-over step) to establish a new BOS. Thus, to maintain lateral stability during the single-limb support phase, the XCOM movement must be restrained within the lateral boundary of the support limb. In contrast, to manoeuvre laterally, the XCOM must ultimately travel beyond the lateral boundary of the support limb.

The inverted pendulum model has two important implications for this study. First, there is an inverse relationship between medial and lateral stability—increasing the MOS in one direction will reduce stability in the other (12,23). Second, gait patterns that increase stability in one direction will impede manoeuvrability in that same direction by passively resisting any self-imposed forces intended to redirect one’s trajectory (8) and ultimately require larger impulses to overcome the body’s resistance to movements (13,24,25). This resistance may increase the time required to perform an effective manoeuvre toward the BOS. Therefore, during periods of single limb support, increasing the lateral MOS will increase stability (resistance to lateral perturbations) but decrease the capacity to manoeuvre quickly in the lateral direction. In addition, increasing the lateral MOS during walking will be beneficial for initiating a medial manoeuvre *away* from the BOS.

The quantification of manoeuvrability was also critical for this study. During legged locomotion, ground reaction forces and impulses are typically used to quantify the magnitude of the control signal that is generate to perform a manoeuvre (26–28). When a direct measure of the control signal is not available, performance metrics are often used as a proxy to gain insight about the magnitude of the control signal and a body’s relative resistance to that control signal. Such measures may include choice reaction time, the number of steps to initiate a manoeuvre, and time to initiate manoeuvre (12,29–31). However, the rate at which a manoeuvre is performed is insufficient to fully characterize the ability to manoeuvre. It is also necessary to evaluate if the actual manoeuvre matches the desired manoeuvre to provide an indication of how appropriate the control signal is. Thus, we chose two proxy measurements to quantify, 1) the body’s general resistance to the manoeuvre control signal and 2) the accuracy of the manoeuvre. These measures were the manoeuvre reaction time (the time required to initiate a manoeuvre), and foot placement error (the distance between the foot and manoeuvre end target), respectively.

Based on our selected methods to quantify gait stability and manoeuvre performance we hypothesized that in the Perturbed environment people would increase their lateral MOS when compared to walking in the Baseline environment. We also hypothesized that in the perturbed environment, people would increase manoeuvre reaction time and foot placement errors for laterally directed manoeuvres but not for medially directed manoeuvres when compared to similar manoeuvres performed in the Baseline environment. Finally, we hypothesized that people would leverage the relationship between stability-manoeuvrability by selecting body states that lower lateral stability when they anticipated having to produce lateral manoeuvres (independent of environment) than when anticipating medial manoeuvres. Our outcomes empirically confirm the mechanical relationship between stability and manoeuvrability during human walking by demonstrating that gait characteristics that improve stability in one direction also result in slower manoeuvres in that direction, but not in the opposite direction. Moreover, we demonstrate that humans leverage the mechanical stability-manoeuvrability relationship by selecting behaviours that change their state along this continuum in a manner that considers both changing task goals (the need to manoeuvre) and external environment (the need to resist perturbations).

## 2.0 Methods

### 2.1 Participants

Twenty-four adults (25.6 ± 2.9 years, height 1.7 ± 0.1 m, weight 72.8 ± 14.2 kg, 14 males, mean ± standard deviation) participated. The Northwestern Institutional Review Board (IRB) approved the protocol, and all participants gave written informed consent. Participants did not have any known musculoskeletal or cardiovascular injuries/impairments affecting their gait or balance. All participants self-reported the ability to walk continuously for 30 minutes without experiencing fatigue, health risks, or other limitations. Twenty-one participants self-reported right-foot dominance.

### 2.2 Experimental Setup

Participants performed a series of discrete stepping trials that involved stepping quickly from a start target to an end target. They were instructed to initiate every trial with their right leg and to try to reach the end target in three steps. During each trial, participants had the chance of experiencing disruptive forces as well as the chance of the target suddenly shifting medially or laterally after starting their movements. Prior to the start of the experiment, participants performed practice trials where they walked both straight and to the shifted target, with and without disruptive forces.

The targets were projected on the floor using a short-throw projector (Hitachi America, ltd.). Each target consisted of a pair of parallel ellipses, one for each foot, with 0.3 m major and 0.15 m minor axes and no gaps between the ellipses (Fig. 1A). The end target location was located 1.5 leg lengths in front of the start target at the start of all trials. To mitigate the risk of falls, participants wore a trunk harness attached to an overhead safety support trolly that allowed free fore-aft motion. A string potentiometer was attached to the front of the harness to monitor participants’ real-time forward walking velocity. An audible “beep” cued the start and end of each trial. To encourage similar walking speeds across participants (1.0-1.3 m/s), peak forward velocity was displayed on a monitor positioned in front of the walking path during practice trials. However, to encourage participants to prioritize accuracy over speed, the feedback was then removed after practice trials and was not provided during the experimental testing.

**Figure 1.**
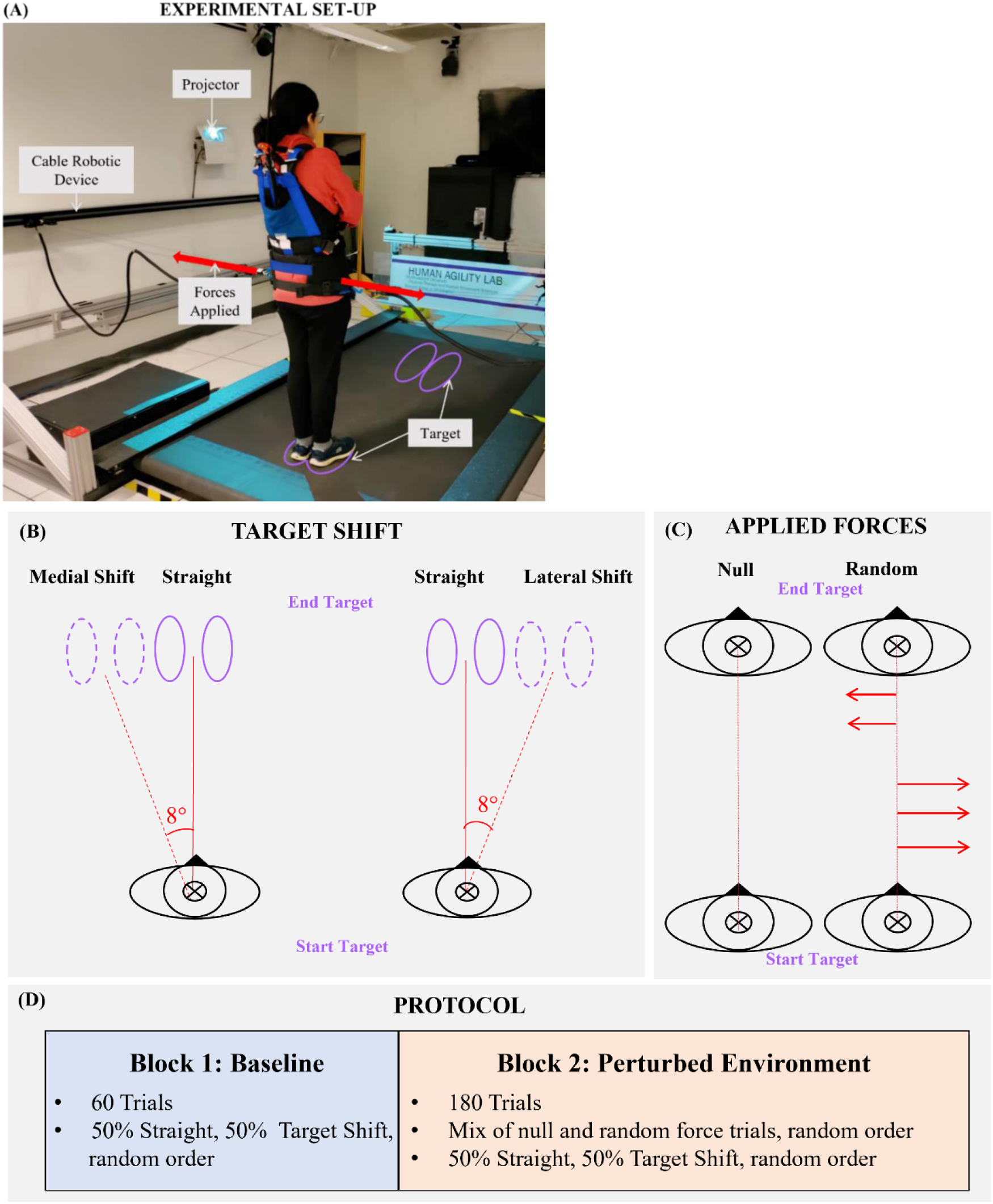
(A) Experimental Set-Up: Participants performed a series of stepping trials. Start and end targets consisted of a pair of parallel ellipses projected using a short-throw projector. The end target was at a distance of 1.5x their leg length. Medio-lateral forces were applied through a pelvic harness worn by the participant using a cable-driven robot on select trials. (B) Target Shift: Representation of a medial and lateral target shift. On select trials, the target was instantaneously moved after the participant began forward walking. Note that this figure is intended as an illustrative sketch, the actual dimensions of the target can be found in section 2.2. (C) Applied Forces: Types of forces applied by the cable robot during select trials (D) Protocol: Participants performed 240 trials, split into two blocks – Baseline and Perturbed environment.

For half of the trials, the end target remained directly in front of the start target (Straight trials). For the other half of trials, the end target shifted mid-trial 8° laterally (Target Shift trials; Fig. 1B). Participants were randomly assigned to either a Lateral or Medial manoeuvre group that would experience target shifts exclusively to the right (Lateral; 12 participants) or left (Medial; 12 participants), respectively, for the entire experiment. During Target Shift trials, the target location would jump to the shifted location after participants had begun moving forward but also prior to (or just during) heel strike of the first step. Pilot testing revealed that a forward velocity threshold of 0.4 m/s accomplished these manipulation goals for all participants and was used as the threshold for triggering the target shift. Thus, participants did not learn of the end target location for each trial until they began walking forward (“go before you know”). This setup encouraged participants to react to the target movement rather than try to pre-plan movement trajectories for each trial.

Forces were applied medio-laterally to the participants’ pelvis using a cable-driven robot (Fig. 1A). The robot consisted of two independent series elastic actuators (linear motor and spring) that transmitted bilateral forces through a cable transmission system to a snug pelvic harness worn by the participant (32). Depending on the trial, a custom-written LabVIEW (National Instruments) code commanded the robot to apply one of two patterns of forces to the participant (Fig. 1C):

> Null Forces: Zero net medio-lateral force applied during the stepping trial.
>
> Random Forces: A series of square-wave impulse forces. Each impulse force had a randomly generated direction (left or right), magnitude (75-90 N), and duration (300-500 ms). The total impulses applied for each trial were randomized to be either 1 or 2. Forces were checked during pilot testing to ensure the vector sum of all total forces applied during a session was 0 N.

We collected motion capture data to estimate the whole-body centre-of-mass (COM) trajectory and measure foot placement location for each trial. We collected kinematic 3D marker data using a 12-camera motion capture system (Qualisys, Gothenburg Sweden) sampling at 100 Hz. We placed 15 passive reflective markers on the pelvis and bilaterally on the greater trochanter, lateral malleolus, calcaneus, and the 2nd, 3rd, and 5th metatarsals. Our outcome measures were the Minimum Lateral Margin of Stability, Manoeuvre Reaction Time, Foot Placement Error, and Step Width and Length (Supplemental Materials).

### 2.3 Protocol

Participants first performed eight practice trials. These provided experience stepping to end targets that were straight ahead or shifted laterally during the trial. They also provided practice of a mix of Null and Random force trials.

Participants then performed a total of 240 discrete stepping trials, consisting of a block of 60 Baseline trials (Null forces every trial), followed by a block of 180 trials in a Perturbed environment (a mix of 120 Random and 60 Null force trials that varied trial-to-trial) (Fig 1D). The order of the Random and Null forces trials was pseudo-randomized such that each set of three trials contained two Random trials and one Null trial, and their order within each triplet was randomized. The target shifted for half of the trials in each block (30 in Baseline; 90 in the Perturbed environment), and did not shift (Straight trials) for the other half. Likewise, the target shifted for half of the Random and half of the Null trials in each block (30 Null trials in Baseline; 30 Null trials and 60 Random trials in the Perturbed environment) and did not shift for the other half. The order of Target Shift and Straight trials was randomized for each block.

For each trial, participants always stepped first with their right leg. Participants were not informed if they would experience Null or Random forces each trial or whether the target would shift location. Participants were instructed to start the trial when they heard a start “beep” and to step to the target as accurately as possible. The trial concluded when the participant had taken 3 steps to the target (the first right step, then a left step that would ideally land in the left end target, and finally another right step that would ideally land in the right end target). A stop “beep” indicated the trial was complete (triggered manually by the experimenter after observing the participant complete the trial). No forces were applied as participants stepped backwards to reset to the start target location for the next trial. To minimize the possible effect of arm swing on stability and manoeuvrability, participants crossed their arms across their chest during the stepping trials. Participants were instructed to step quickly and accurately to the end target with their arms crossed, and to walk in a manner that they felt was comfortable while complying with the primary objective. No further instructions were given.

### 2.4 Data Processing

Kinematic data was processed using Visual3D (C-Motion, Germantown, MD) and a custom-written MATLAB (MathWorks, Natick, MA) program. Marker data was gap-filled and passed through a low-pass Butterworth filter with a cut-off frequency of 6 Hz. The position of the COM was estimated using Visual3D as the centre of a pelvis model created from the pelvis markers (33) and the bilateral greater trochanter markers. Heel-strike and toe-off gait events were identified in MATLAB using the calcaneus and 2nd metatarsal markers, respectively. Toe-off was identified as the local minimum in the vertical position of the 2nd metatarsal marker just prior to the maximum position associated with the swing phase of the foot. Heel strike was then identified as the local minimum in the vertical position of the calcaneus marker just following this maximum. All gait events were visually inspected to verify event detection and corrected as necessary.

### 2.5 Data Analysis

#### 2.51 Lateral Margin of Stability (MOS)

We quantified lateral stability using the minimum lateral MOS. We calculated the minimum lateral MOS for the stance phase of the first right step. We were interested in assessing systemic adaptations to stability – general adaptations implemented regardless of target shift or perturbations. Thus, to avoid MOS that could be a combination of systemic adaptations and reactions to target shift/perturbations, we assessed MOS for trials when the target did not shift location (Straight trials) and only Null forces were applied (30 Baseline environment trials, 30 Perturbed environment trials) (Fig. 2A).

**Figure 2.**
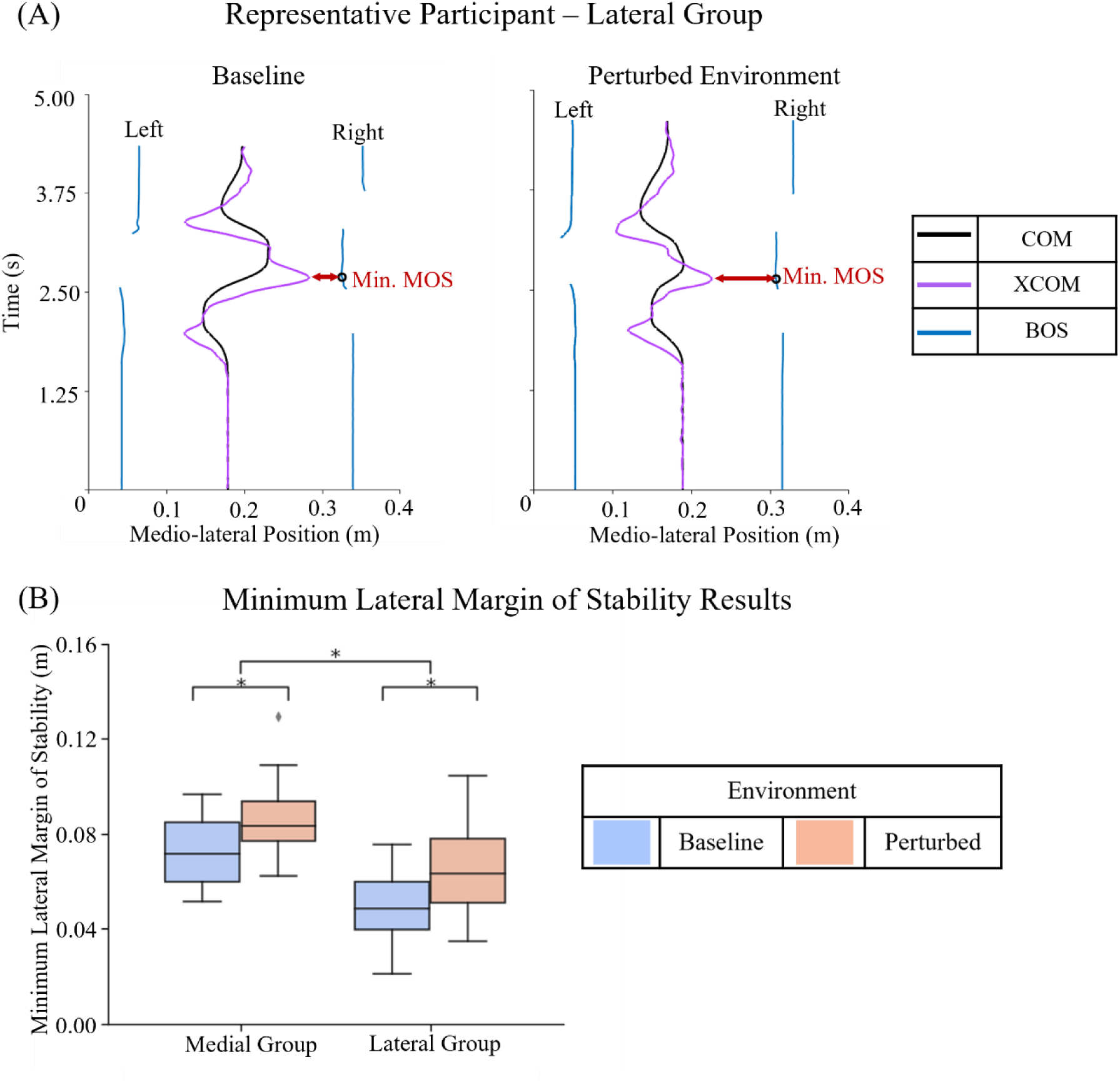
Lateral Margin of Stability (A) Representative Participant: Minimum lateral margin of stability calculation during selected trials during baseline and the Perturbed environment. Note that for specific Perturbed environment trial, no external medio-lateral forces were applied to the participant. The arrow in red represents the minimum margin of stability for the first right step (Eq. 1) (B) Group Data: Box plots and statistical results for minimum lateral margin of stability. Significance (p<0.05) is denoted by *.

The minimum lateral MOS is a function of the extrapolated COM (XCOM) and the lateral edge of the base of support (BOS), as defined in Eq (1):

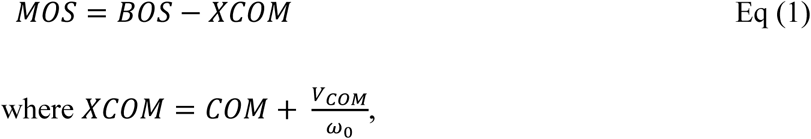

where V is the lateral COM velocity and 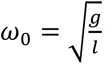 where *g* is the acceleration due to gravity and *l* is the pendulum length (leg length) of the participant (19). The lateral boundary of the BOS was defined as the marker on the 5^th^ metatarsal of the right foot.

The magnitude of a lateral impulse required to move the XCOM beyond the lateral BOS increases proportionately with the minimum lateral MOS. Thus, increases in the minimum lateral MOS indicate increased resistance to lateral impulses and, inversely, decreased resistance to medial impulses.

#### 2.52 Manoeuvre Reaction Time

We were interested in how manoeuvrability was affected by behavioural adaptations accommodating the task goal (making unexpected manoeuvres) and environment. To eliminate confounds from the perturbations themselves, we analysed only trials during which Null forces were applied and the target shifted (30 Baseline, 30 Perturbed Environment).

As a measure of manoeuvrability, we assessed how quickly participants were able to carry out the manoeuvre after reacting to each target shift. While previous research has used a choice reaction time measure for stepping (29,30) or time and number of steps before a manoeuvre for continuous walking tasks (12,31), we sought a different approach to account for the fact that participants were already moving and were restricted to three steps when tasked with reacting to the target shift. Manoeuvre reaction time (*τ_reaction time_*) was defined statistically as the time when the participants’ manoeuvre trajectory in a Target Shift trial first differed from a Straight trial (Fig 3A). To make this estimate, we first calculated a 95% confidence interval (CI) of the COM trajectories for Straight trials that occurred with Null forces in both the Baseline and Perturbed environments. Then, for the COM trajectory of each Target Shift trial in each environment, a deviation time ( *τ_deviation_*) was defined as the time when the COM trajectory moved outside of that environment’s Straight trial CI in the same direction as the end target shift. If no deviation occurred, the time of deviation was assigned a value equal to the time required to complete the trial. To standardize the manoeuvre reaction time across each trial and participant, we used the time of target shift (*τ*_*target shift*_) as the cue for participants to begin the manoeuvre. Thus, manoeuvre reaction time for each trial was calculated as shown in Eq (2).

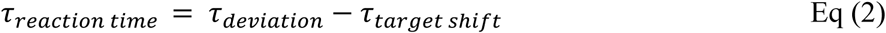

**Figure 3.**
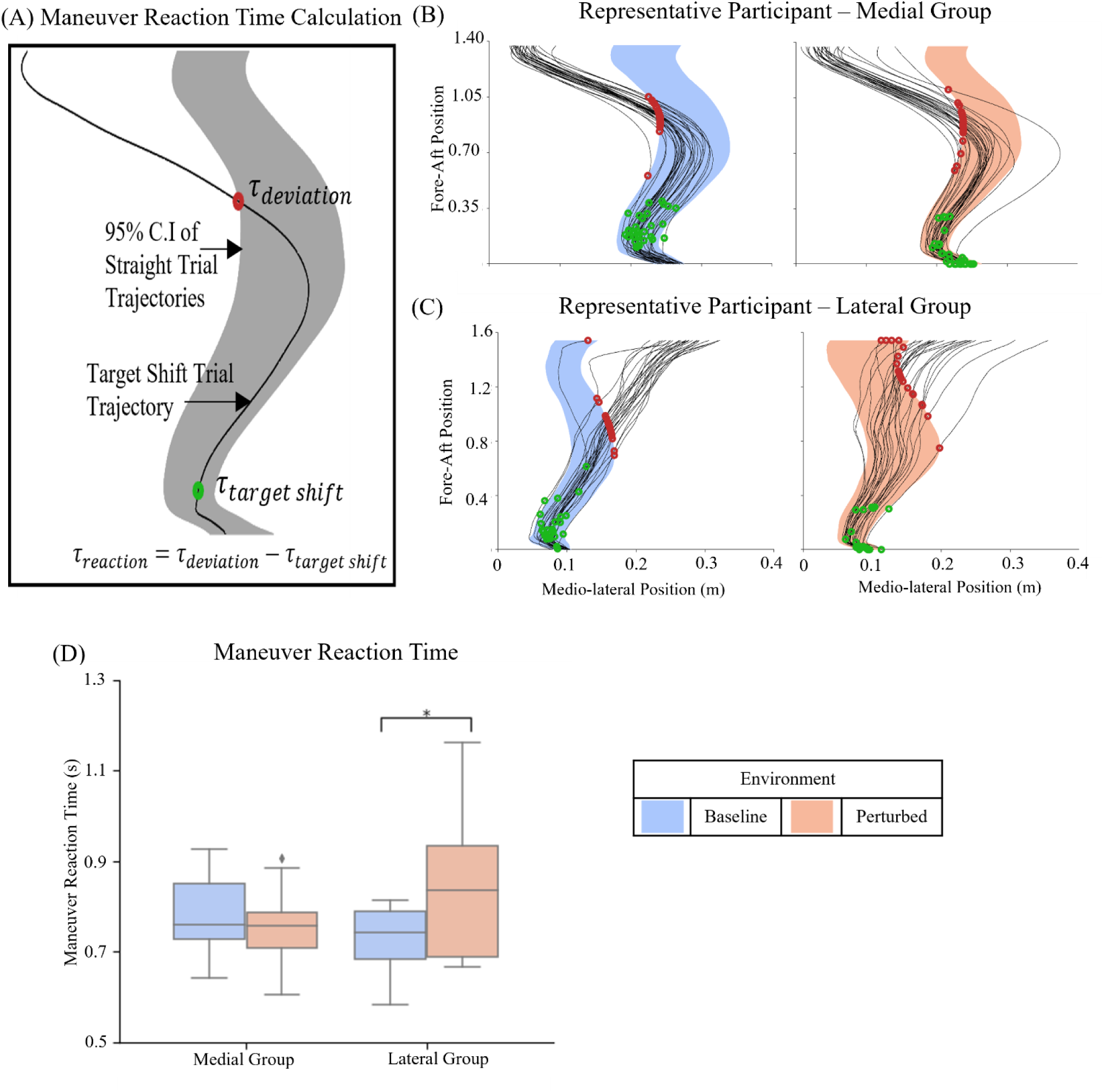
Manoeuvre Reaction Time (A) Reaction Time Calculation: Calculation of reaction time for a representative participant for a single medial Target Shift trial. The shaded region represents the 95% confidence interval of all Straight-walking COM trajectories in either Baseline or the Perturbed environment. The black line represents the COM trajectory for a single Target Shift trial for a participant in the Medial group. (B) Representative Participant: COM trajectories for all Target Shift trials for a representative participant from the Lateral group. The green dots indicate the moment the target shifted for a trial. The red dots represent the point of deviation of a Target Shift trial COM trajectory from the confidence interval of Straight-walking trials (C) Representative Participant: COM trajectories for all Target Shift trials for a representative participant from the Medial group. (D) Group Data: Box plots and statistical results for manoeuvre reaction time. Significance (p<0.05) is denoted by *.

#### 2.53 Foot Placement Error

To further understand participants’ systemic adaptations of manoeuvrability to both environment (Baseline or Perturbed) and the effect of manoeuvre direction (Lateral or Medial), we examined foot placement error. We measured how accurately participants placed their feet within the end targets each trial. Specifically, we calculated a cumulative foot placement error value from the relative final locations of both the left and right foot relative to the left and right end target locations (Fig 4A). Like manoeuvre reaction time, we analysed only trials where Null forces were applied and the end target shifted (30 Baseline environment trials, 30 Perturbed environment trials). Foot placement error was determined as the Euclidean distance between the centre of the foot and the centre of the respective end target. The centre of the foot was defined as the centroid of a polygon formed using the markers on the 2nd and 5th metatarsals and the calcaneus. The centre of the target was the centroid of the elliptical target for each foot. The error for a trial was defined as the mean of the Euclidean distances for both feet. Successful trials were defined as trials where error was less than 0.15 m (the maximum Euclidean distance from the centre of each target to the target boundary).

**Figure 4.**
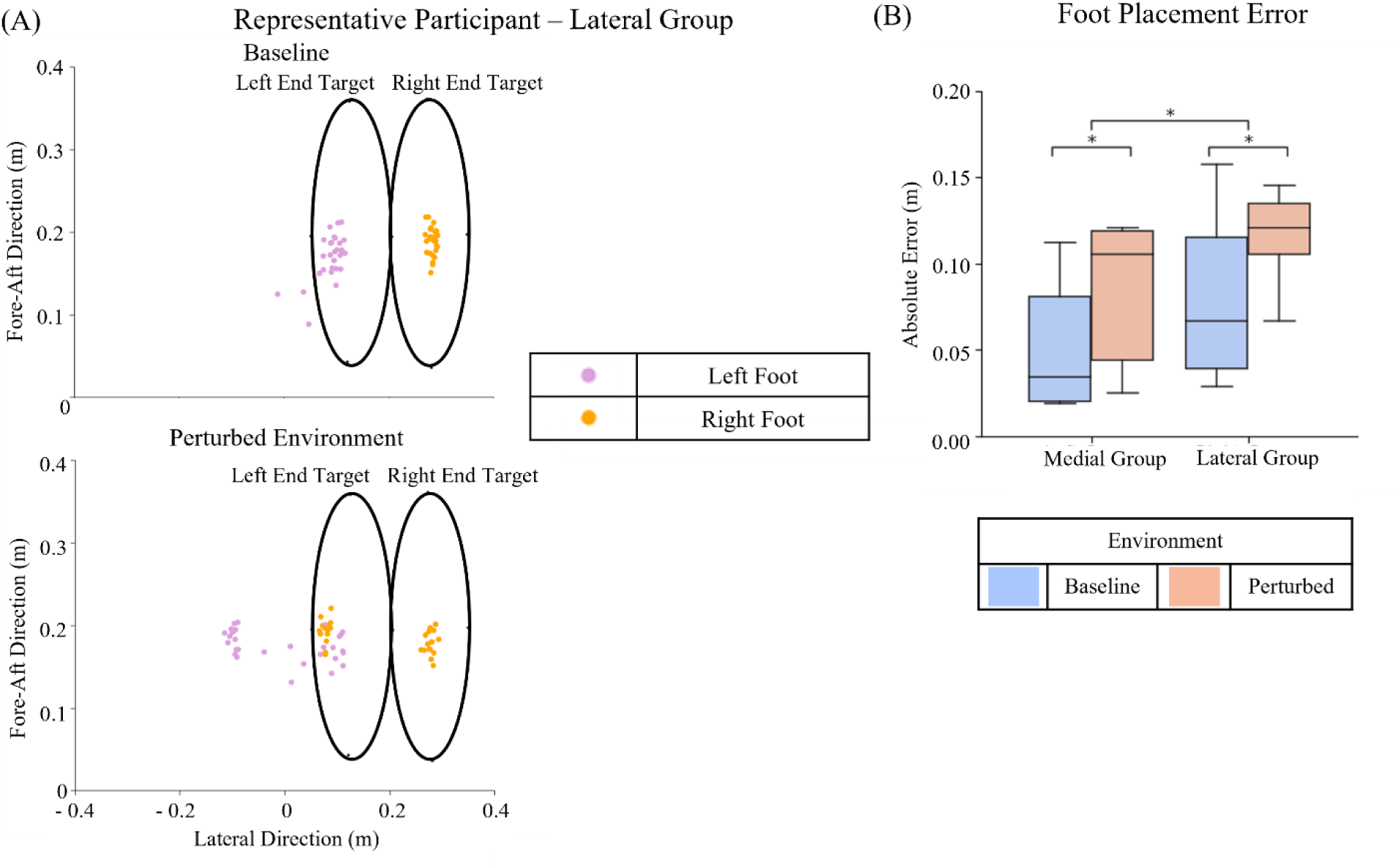
Foot Placement Error (A) Representative Participant: Foot placement for all Target Shift trials for a representative participant from the Lateral group. Here the 0 m coordinate on the x-axis denotes the lateral midpoint of the target for Straight trials. (B) Group Data: Box plots and statistical results for foot placement error. Significance (p<0.05) is denoted by *.

### 2.6 Statistical Analysis

The number of participants was selected based on an *a priori* power analysis conducted assuming repeated-measures ANOVA examining within-between interactions. We used an effect size of 0.48 obtained from pilot trials, 95% power, and two groups with two factors. The power analysis indicated that 9 participants in each group were required. We chose a slightly larger sample size than that calculated in the power analysis to be conservative as the effect size chosen was based on a small sample size.

As the application of mediolateral forces during the Perturbed environment can directly influence a manoeuvre, only trials where Null forces were applied were processed and analysed. To evaluate the effect of the environment, Null force trials in the Baseline and Perturbed environments were compared.

To detect differences in the minimum lateral MOS between Baseline and Perturbed environments, we calculated an average value for each participant from the 30 Straight trials performed during Baseline and from the 30 Straight trials performed during Null trials in the Perturbed environment. To detect differences in the manoeuvre reaction time between Baseline and Perturbed environments, we calculated an average value for each participant from the 30 Target Shift trials performed during Baseline and from the 30 Target Shift trials performed during Null trials in the Perturbed environment. To detect differences in the foot placement error between Baseline and Perturbed environment, we calculated an average value for each participant from the 30 Target Shift trials performed during Baseline and the 30 Target Shift trials in the Perturbed environment. The data were not normally distributed, and therefore non-parametric tests were used. We performed Kruskal tests with a between factor of group (Medial or Lateral) and Friedman tests with a within factor of environment (Baseline or Complex). When a significant main effect of environment was found, pairwise Wilcoxon tests were conducted.

All statistical analyses were conducted using the Pingouin package in Python (34). Normality of data was checked using the Shapiro-Wilkes test. Significance was set at p<0.05 for all analyses. When multiple comparisons were made, a Bonferroni correction was applied. We performed a repeated-measures mixed ANOVA with a between factor of group (Medial or Lateral manoeuvre) and within factor of environment (Baseline or Perturbed) to evaluate differences. When a significant main effect of environment was found, pairwise t-tests comparisons were made.

## 3.0 Results

All 24 enrolled participants were able to successfully complete the full experimental protocol. All participants’ data were included in the analyses.

### 3.1 Lateral Margin of Stability

The minimum lateral MOS was significantly larger in the Perturbed environment than Baseline for both groups (p<0.001, ANOVA) and was larger for the Medial group than the Lateral group (p=0.0024, ANOVA) (Fig 2B). Minimum lateral MOS increased from 0.072±0.014 m during Baseline to 0.086±0.017 m during Perturbed environment trials for the Medial group (Bonferroni corrected-p=0.0070, Paired t-test). Minimum lateral MOS also increased from 0.048±0.015 m during Baseline to 0.06±0.019 m during Perturbed environment trials for the Lateral group (Bonferroni corrected-p=0.0035, Paired t-test).

### 3.2 Manoeuvre Reaction Time

The target shifted prior to heel strike of the first step (153.05±107.76 ms prior to heel strike) for nearly all (91.56%) of the target shift trials, confirming that our manipulation goal to have the target shift prior to (or just during) heel strike was successful.

Statistical analysis revealed a significant main effect for the environment (p=0.048, ANOVA) but not group (p=0.80, ANOVA) (Fig. 3D). An interaction effect (p=0.0045, ANOVA) was found between environment and group. The paired t-tests found significantly delayed manoeuvre reaction times in the Perturbed environment than Baseline when manoeuvring laterally (Bonferroni corrected-p=0.023, Paired t-test). There were no significant differences in manoeuvre reaction time between the Perturbed environment and Baseline when manoeuvring medially (Bonferroni corrected-p=0.59, Paired t-test).

Manoeuvre reaction time increased from 0.73±0.075 s during Baseline trials to 0.83±0.14 s in Perturbed environment for the Lateral group. Manoeuvre reaction time was 0.78±0.083 s during Baseline trials and 0.76±0.078 s in Perturbed environment for the Medial group.

### 3.3 Foot Placement Error

Statistical analyses revealed that foot placement was less accurate in the Perturbed environment than Baseline (p<0.001, Friedman) and was less accurate for the Lateral group than the Medial group (p=0.0054, Kruskal) (Fig. 4B). The distance between the centre of the foot and target increased from 0.08±0.044 m during Baseline to 0.11±0.023 m in the Perturbed environment for the Lateral group (Bonferroni corrected-p =0.013, Wilcoxon). The distance between the foot and target also increased from 0.049±0.032 m during Baseline to 0.082±0.038 m in the Perturbed environment for the Medial group (Bonferroni corrected-p <0.001, Wilcoxon). Successful trials made up 70.26% of total trials. For the Lateral group, 73.59% of Baseline trials and 60.58% of Perturbed environment trials were successful. While for the Medial group, 82.92% of Baseline trials and 64.58% of Perturbed environment trials were successful.

## 4.0 Discussion

We investigated how gait adaptations that increase lateral stability affect the ability to perform laterally and medially directed manoeuvres during walking. We hypothesized that increasing the lateral MOS—a common gait adaptation for increasing lateral stability—would impair the ability to make lateral manoeuvres but facilitate medial manoeuvres. Our findings support our hypothesis—participants increased lateral MOS in the Perturbed environment when compared with the Baseline environment. This increase in lateral MOS coincided with an increase in manoeuvre reaction time for laterally but not medially directed manoeuvres. This outcome confirms the known mechanical stability-manoeuvrability relationship: increasing stability in the lateral direction impedes lateral manoeuvres but facilitates medial manoeuvres. In addition, we observed that people selected a smaller lateral MOS when they anticipated making a lateral manoeuvre when compared to people anticipating medial manoeuvres. This modulation of the lateral MOS in anticipation of walking task was observed in both the Baseline and Perturbed environments. These results indicate that people make behavioural modification to their underlying gait patterns to favourably exploit the mechanical relationship between lateral stability and manoeuvrability depending on anticipated needs to manoeuvre and/or resist external perturbations.

### 4.1 Lateral Margin of Stability

We found participants increased their minimum lateral MOS in response to two factors; the possibility of receiving a medio-lateral perturbation, and the possibility of having to initiate a medially directed manoeuvre.

All participants increased their minimum lateral MOS when performing the stepping task in the Perturbed environment when compared to Baseline. Increasing the minimum lateral MOS in the Perturbed environment increases the body’s passive resistance to lateral perturbations (19,20). This occurs because the gravitational moment about the ankle-joint in the frontal plane that acts to accelerate the COM medially increases as the distance between the XCOM and the lateral base of support increases (35). Our observation that people increased their lateral MOS in response to lateral perturbations was consistent with several prior studies that also observed increases in lateral MOS when people walked in environments where they were exposed to unpredictable lateral perturbations (9,10). This outcome indicates that people make behavioural modifications to their underlying gait patterns to bias the stability-manoeuvrability relationship towards greater lateral stability in environments where there is threat to frontal-plane stability.

We also found the group required to make Medial manoeuvres had a significantly larger lateral MOS during the first step of the task (always the right limb in stance phase) than the group required to make Lateral manoeuvres. The effect was observable even in the Baseline environment. This demonstrates that people make behavioural modifications to their underlying lateral MOS directly in response to the potential need to initiate a medial or laterally directed manoeuvre. Increasing the minimum lateral MOS acts to accelerate the COM medially (away from the stance limb) (35). An increased lateral MOS on the right limb should be beneficial for initiating a medial manoeuvre because this body position will decrease medial stability and thereby reduce any passive resistance to medial movements. In addition, when compared to a foot placed directly below the pelvis, placing the right limb lateral to the body allows the relatively large extensor muscles of the hip, knee, and ankle joints to produce lateral directed ground reaction force (GRF) to assist with the manoeuvre. While not measured directly in our study, others have discussed this as a possible method to facilitate performance of lateral manoeuvres (23).

During manoeuvre trials we chose to analyse manoeuvre reaction time during the stance phase of the first step. We performed a post-hoc analysis to confirm how frequently manoeuvres also began during the stance phase of the first step. We found that for all Target Shift trials (30 Baseline and 30 Perturbed) across all 24 participants, the deviation time was 0.126 ± 0.148 s before the toe-off of the first right step. The deviation time occurred during the stance phase of the first step for 81.4% (1139 out of 1399) of trials where a manoeuvre occurred (trials with no deviation point, i.e. the reaction time was the total trial time, were excluded from this analysis). Thus, for a large majority of trials, the margin of stability calculated during the stance phase of the first step is representative of the strategy adopted by participants in anticipation of both the external challenge to stability and the need to manoeuvre.

In this study, we quantified lateral MOS during the first step of the task in each environment (Baseline and Perturbed) only during trials when both the target did not shift position and only null forces were applied. As participants were 1) not informed of target shifts or the forces they would experience and 2) the onset of both target shifts and random forces did not begin until forward walking was initiated, we believe that the lateral MOS during the first step is representative of a stepping strategy participants adapted in anticipation of the environment. However, it is possible that participants may have been able to modulate their lateral MOS after beginning forward stepping if they realized that the target was not going to shift, or that random forces would not be applied. If this were the case, the effect would be to decrease the lateral MOS which contrasts with our observation that lateral MOS increased. Thus, our major conclusion that MOS increased in the Perturbed environment is unlikely to have been affected.

### 4.2 Manoeuvrability (Manoeuvre Reaction Time and Foot Placement Error)

We observed a significant increase in manoeuvre reaction time in the Perturbed environment when compared to Baseline for lateral manoeuvres. In contrast, we found no significant difference in manoeuvre reaction time between the Perturbed environment and Baseline for medial manoeuvres. Additionally, we observed significantly higher foot placement error in the Perturbed environment for both groups. We also found significantly lower foot placement error in the Medial group compared to the Lateral group, even at Baseline.

As described earlier, the manoeuvre reaction time outcomes can be explained by the increase in lateral MOS observed in the Perturbed environment. For the Lateral group, an increase in lateral MOS of the right limb in the Perturbed environment resulted in a longer manoeuvre reaction time and greater foot placement error because this body position resisted the participants’ volitional efforts to manoeuvre laterally, leading to a stability-manoeuvrability trade-off.

In contrast, we did not observe a stability-manoeuvrability trade-off for the Medial group. The increase in lateral MOS entails a decrease in the medial MOS, which resulted in a body state that reduced the resistance to the participants’ volitional efforts to manoeuvre medially. Furthermore, while the foot placement error increased in the Perturbed environment, it was significantly lower than the Lateral group in both Baseline and the Perturbed environments. Thus, for the Medial group, increasing the lateral MOS on their right limb resulted in a body position that was favourable for both resisting lateral perturbations and making medial manoeuvres. Therefore, our findings confirm this directionality in the stability-manoeuvrability trade-off during walking due to the inverse relation between lateral and medial MOS.

There are other factors in addition to lateral MOS that may have decreased manoeuvrability for Lateral manoeuvres but not for Medial manoeuvres. One explanation for this observed directionality may be the inherent anthropometric challenges of performing a laterally-directed manoeuvre. Participants may have found it more difficult to perform a “cross-over” step to manoeuvre to a laterally shifted target than to perform a side-step to manoeuvre to a medially shifted target. Previous research suggests that people struggle to initiate manoeuvres in stepping tasks when the signal to manoeuvre is given on a step that is on the same side as the manoeuvre direction (31). Variability in the time of target shift relative to first right heel strike may have also affected foot placement error. However, while a post-hoc correlation analysis between time of target shift relative to the first right heel strike revealed a significant correlation (Spearman, p<0.01), the variability in the foot placement error captured by this correlation is very small (R^2^ = 0.0116). Therefore, while a later target shift may have led to slightly higher foot placement error, it cannot explain our results. Prior research has also shown larger absolute errors in stepping tasks where the targets are on the opposite side of the stepping foot (36). Thus, for this paradigm, a target shift to the right (Lateral group) may have been more challenging as the first foot entering the target was the participants’ left foot. Therefore, the Medial group may have found it less challenging to manoeuvre during a Target Shift trial than the Lateral group.

### 4.3 Limitations

While our results provide indirect support of our hypothesis—increases in lateral MOS occurred in the same conditions that indicated increases in lateral (but not medial) manoeuvre reaction time—we were not able to formally demonstrate the principle by directly associating MOS values with manoeuvre reaction times. As mentioned in the analysis sections (2.51 and 2.52), we assessed lateral MOS in Straight trials but assessed manoeuvre reaction time in Target Shift trials. While this was done so that lateral MOS could be assessed without the possible confounds of volitional reactions to the target shift, it precluded us from making a direct comparison between values. In addition, one of our measures of manoeuvrability – foot placement error – always occurred during the step(s) after our assessment of stability was made. As such, foot placement accuracy will be the result of both the appropriateness of the initial manoeuvre control signal to create the desired movement trajectory and any corrective actions occurring during the subsequent step. While our results find a significant decrease in foot placement accuracy during trials performed in the Perturbation environment than those preformed during Baseline, it is possible that factors other than changes in MOS may have contributed to this change.

## 5.0 Conclusion

The present study confirmed the mechanical relationship between stability and manoeuvrability during human walking by demonstrating that increasing frontal-plane stability results in slower manoeuvres in a given direction. Importantly, the study also demonstrates that people make behavioural changes to their underlying walking patterns that bias their state along the mechanical stability-manoeuvrability relationship in a manner that considers both changing task goals (the need to manoeuvre) and external environment (the need to resist perturbations). Thus, stability and manoeuvrability during human walking results from a dynamic interplay between mechanical and behavioural factors.

## 7.0 Acknowledgements

We would like to thank Jordan Dembsky for his aid in data collection and data processing and the members of the Human Agility Lab for their valued feedback.

## 9.0 Data Availability Statement

The data are available at https://doi.org/10.21985/n2-g778-5k68.

## 10.0 Preprint

A preprint of the article has been published (37).

