## Supplemental Materials for "Humans Exploit the Trade-Off Between Lateral Stability and Manoeuvrability During Walking"

To further understand participants' systemic adaptations of stability and manoeuvrability to both environment (Baseline or Perturbed) and the effect of manoeuvre direction (Lateral or Medial), we examined step width and step length.

### METHODS:

We calculated step width and step length for the first right step on each trial to gain a better understanding of the strategies people were using to generate stability and manoeuvrability. Both metrics were calculated using the location of the centre of the foot at the start of the trial and the location of the centre of the foot at the time point of their minimum lateral MOS. Step width was defined as the mediolateral distance between the centre of the foot at the start of the trial and during stance phase and step length was the fore-aft distance for the same. Step width and step length values were normalized to leg length for each participant.

### STATISTICAL METHODS:

To detect differences in the foot placement error between Baseline and Perturbed environment, we calculated an average value for each participant from the 30 Target Shift trials performed during Baseline and the 30 Target Shift trials in the Perturbed environment. The data were not normally distributed, and therefore non-parametric tests were used. We performed Kruskal tests with a between factor of group (Medial or Lateral) and Friedman tests with a within factor of environment (Baseline or Complex). When a significant main effect of environment was found, pairwise Wilcoxon tests were conducted.

### RESULTS:

Statistical analyses did not find significant differences in step width between stepping trials in either environment ($p=0.68$, Friedman), or between groups ($p=0.099$, Kruskal).

Step length was longer in the Lateral group than Medial. For the Lateral Group step length was longer during Baseline than in the Perturbed environment. We found a significant difference between group ($p<0.001$, Kruskal) and within environment ($p=0.041$, Friedman) for step length. The normalized step length decreased from 0.86±0.056 leg lengths during Baseline to 0.82±0.054 leg lengths in the Perturbed environment for the Lateral group (Bonferroni corrected-$p<0.001$, Wilcoxon) and was 0.77±0.0323 leg lengths during Baseline and 0.77±0.043 leg lengths in the Perturbed environment for the Medial group (Bonferroni corrected-$p =1$, Wilcoxon).
